# Phylogenomics revealed migration routes and adaptive radiation timing of Holarctic malaria vectors of the Maculipennis group

**DOI:** 10.1101/2022.08.10.503503

**Authors:** Andrey A. Yurchenko, Anastasia N. Naumenko, Gleb N. Artemov, Dmitry A. Karagodin, James M. Hodge, Alena I. Velichevskaya, Alina A. Kokhanenko, Semen M. Bondarenko, Mohammad R. Abai, Maryam Kamali, Mikhail I. Gordeev, Anton V. Moskaev, Beniamino Caputo, Sargis A. Aghayan, Elina M. Baricheva, Vladimir N. Stegniy, Maria V. Sharakhova, Igor V. Sharakhov

## Abstract

**Background:** Understanding the evolutionary relationships between closely related taxa is important for mosquitoes that transmit human diseases. Six out of 41 dominant malaria vectors in the world belong to the Maculipennis group, which is subdivided into two North American subgroups (Freeborni and Quadrimaculatus), and one Eurasian (Maculipennis) subgroup. Although previous studies considered the Nearctic subgroups as ancestral, details about their relationship with the Palearctic subgroup, and their migration times and routes from North America to Eurasia remain controversial. The Eurasian species *An. beklemishevi* is currently included in the North American Quadrimaculatus subgroup adding to the uncertainties in mosquito systematics.

**Results:** To reconstruct historic relationships between the North American and Eurasian mosquitoes, we conducted a phylogenomic analysis of 11 Palearctic and 2 Nearctic species based on 1271 orthologous genes using their transcriptomic or genomic sequences. The analysis indicated that the Palearctic species *An. beklemishevi* clusters together with other Eurasian species and represents a basal lineage among them. Also, *An. beklemishevi* is related more closely to *An. freeborni,* which inhabits the Western United States, rather than to *An. quadrimaculatus,* a species from the Eastern United States. The time-calibrated tree suggests a migration of mosquitoes in the Maculipennis group from North America to Eurasia about 20-25 million years ago through the Bering Land Bridge. A Hybridcheck analysis demonstrated highly significant signatures of introgression events between allopatric species *An. labranchiae* and *An. beklemishevi*. The analysis also identified ancestral introgression events between *An. sacharovi* and its Nearctic relative *An. freeborni* despite their current geographic isolation.

**Conclusions:** Our phylogenomic analyses reveal migration routes and adaptive radiation timing of Holarctic malaria vectors and strongly support inclusion of *An. beklemishevi* into the Maculipennis subgroup. The vectorial capacity and the ability to diapause during winter evolved multiple times in Maculipennis evolution. Detailed knowledge of the evolutionary history in the Maculipennis subgroup will help us better understand the current and future patterns of malaria transmission in Eurasia.

## Introduction

The genomics era offers new opportunities for a better understanding of the evolutionary history of species. Unlike traditional molecular phylogenetics, which relies on comparison of only a few sequenced markers, phylogenomics employs a large amount of genomic data based on thousands of Single Nucleotide Polymorphisms (SNP) in the orthologous gene sequences. This feature makes it a powerful and accurate tool for studying evolutionary relationships among groups of organisms involved in the transmission of human pathogens, such as mosquitoes [1–5]. Phylogenomics helps scientists better understand the evolution of epidemiologically important phenotypes including the ability to transmit pathogens [6] and adaptation to the changing natural environment [7, 8]. For example, a comparison of 16 malaria mosquito genomes from different taxonomic groups demonstrated that the ability of mosquitoes to transmit malaria parasites evolved multiple times [9].

The Maculipennis group of malaria mosquitoes has the Holarctic distribution, which covers North America, Europe, Asia, and North Africa [10, 11]. According to the modern classification, the Maculipennis group is subdivided into two North American subgroups, the Freeborni and Quadrimaculatus subgroup, and one Eurasian Maculipennis subgroup [10]. The group belongs to the subgenus Anopheles, which is the only cosmopolitan subgenus among anophelines [12]. The Maculipennis group consists of 11 currently recognized Palearctic species: *Anopheles artemievi,* Gordeyev, Zvantsov, Goryacheva, Shaikevich & Yezhov, 2005; *An. atroparvus,* Van Thiel, 1972; *An. beklemishevi,* Stegniy & Kabanova, 1976; *An. daciae,* Linton, Nicolescu & Harbach, 2004; *An. labranchiae,* Falleroni 1926; *An. maculipennis,* Meigen, 1818; *An. martinius,* Shingarev, 1926; *An. melanoon,* Hackett, 1934; *An. messeae* Falleroni, 1926; *An. persiensis,* Linton, Sedaghat & Harbach, 2003; and *An. sacharovi* Favre, 1903. Six out of 41 dominant malaria vectors in the world belong to the Maculipennis group: *An. atroparvus*, *An. labranchiae*, *An. messeae,* and *An. sacharovi* in Eurasia, as well as *An. freeborni,* Aitken, 1939 and *An. quadrimaculatus,* Say, 1824 in North America [13, 14]. Systematics studies of the Maculipennis group, the former *Anopheles maculipennis* complex, have a long history. Originally, the phenomenon of “anophelism without malaria” in Europe led to the conclusion that *An. maculipennis* represents a complex of species with different abilities to transmit malaria [15, 16]. Later, J. Kitzmiller introduced the idea that the origin of the Maculipennis complex in the American tropics was followed by a migration of these mosquitoes from America to Eurasia with further radiation, producing the Palearctic group of species [17]. This idea was based on results from interspecies hybridization within and between the Palearctic and Nearctic members of the group. Studies of reproductive isolation demonstrated that hybridization between Palearctic members produced more viable and fertile progeny than hybridization between Nearctic members, suggesting that the Palearctic species radiated more recently [17–19]. The presence of fertile F1 females in some inter-species crosses points to the possibility of genetic introgression among the members of the Maculipennis group. However, details about the phylogenetic relationships between and within these subgroups, a possibility of inter-species gene flow, and the routes and times of their migration from North America to Eurasia remain uncertain.

The taxonomy of the Maculipennis group based on morphology is very difficult as its members have identical larval and adult characteristics and some of them can be identified only by the differences in the structure of the eggs [20]. The use of cytogenetics methods significantly contributed to species identification. Analyses of polytene and mitotic chromosomes described species-specific features of the mosquito karyotypes such as fixed inversions and heterochromatin structure [17, 21–25]. Cytogenetic photomaps for Palearctic species (*An. atroparvus*, *An. beklemishevi, An. labranchiae, An. maculipennis*, *An. martinius, An. melanoon, An. messeae,* and *An. sacharovi*) have been used to identify fixed overlapping chromosomal inversions [21]. These inversions were employed for the reconstruction of phylogenetic relationships among the members of the Maculipennis subgroup [26]. Based on the assumption of the monophyletic origin of inversions, the Eurasian mosquitoes were divided into three major clades: the European clade (*An. labranchiae-An. atroparvus*), the Northern Eurasian clade (*An. melanoon-An. maculipennis-An. messeae*), and the Southern Eurasian clade (*An. sacharovi-An. martinius*) [21]. Following the principles of phylogeny reconstruction developed by Hennig [27], V.N. Stegniy concluded that if several species are identical in the structure of any chromosomal rearrangements, then that structure is phylogenetically ancestral in comparison with any other unique inversion form. Accordingly, the European clade was defined as the basal clade that gave rise to the Northern Eurasian and Southern Eurasian clades [21, 28]. The Eurasian species *An. beklemishevi* was not included in any of these clades since its chromosomal banding pattern was quite distinct. Although a cytogenetic analysis failed to establish phylogenetic relationships between *An. beklemishevi* and the other Palearctic members, similarities in the banding patterns were noticed between *An. beklemishevi* [21, 22] and the North American species *An. earlei*, Vargas, 1943 [29] from the Freeborni subgroup. The cytogenetic observations, the geographic distribution of the clades [21, 28], and the results of inter-species hybridization [17–19] led V.N. Stegniy to the hypothesis that two independent radiation events of malaria mosquitoes occurred from North America to Eurasia [26] (**Fig. 1**). According to this hypothesis, the first migration of an ancestral species related to the eastern North American species *An. quadrimaculatus* occurred through the Greenland connection between Europe and eastern North America, which existed until the end of the Eocene, ∼50 million years ago (Mya) [30]. This migration resulted in the origin of the European clade that further radiated into the Northern Eurasian and the Southern Eurasian clades. The second migration of an ancestral species related to the western North American subgroup, *An. freeborni*, occurred through the Bering Land Bridge, which existed between Asia and western North America ∼20 Mya [30]. This migration gave rise to *An. beklemishevi.* An ecological study provided indirect support to the idea that *An. beklemishevi*, or its immediate ancestor, migrated into Eurasia *via* the Bering Land Bridge where the species’ range underwent substantial expansion [31].

**Figure 1.**
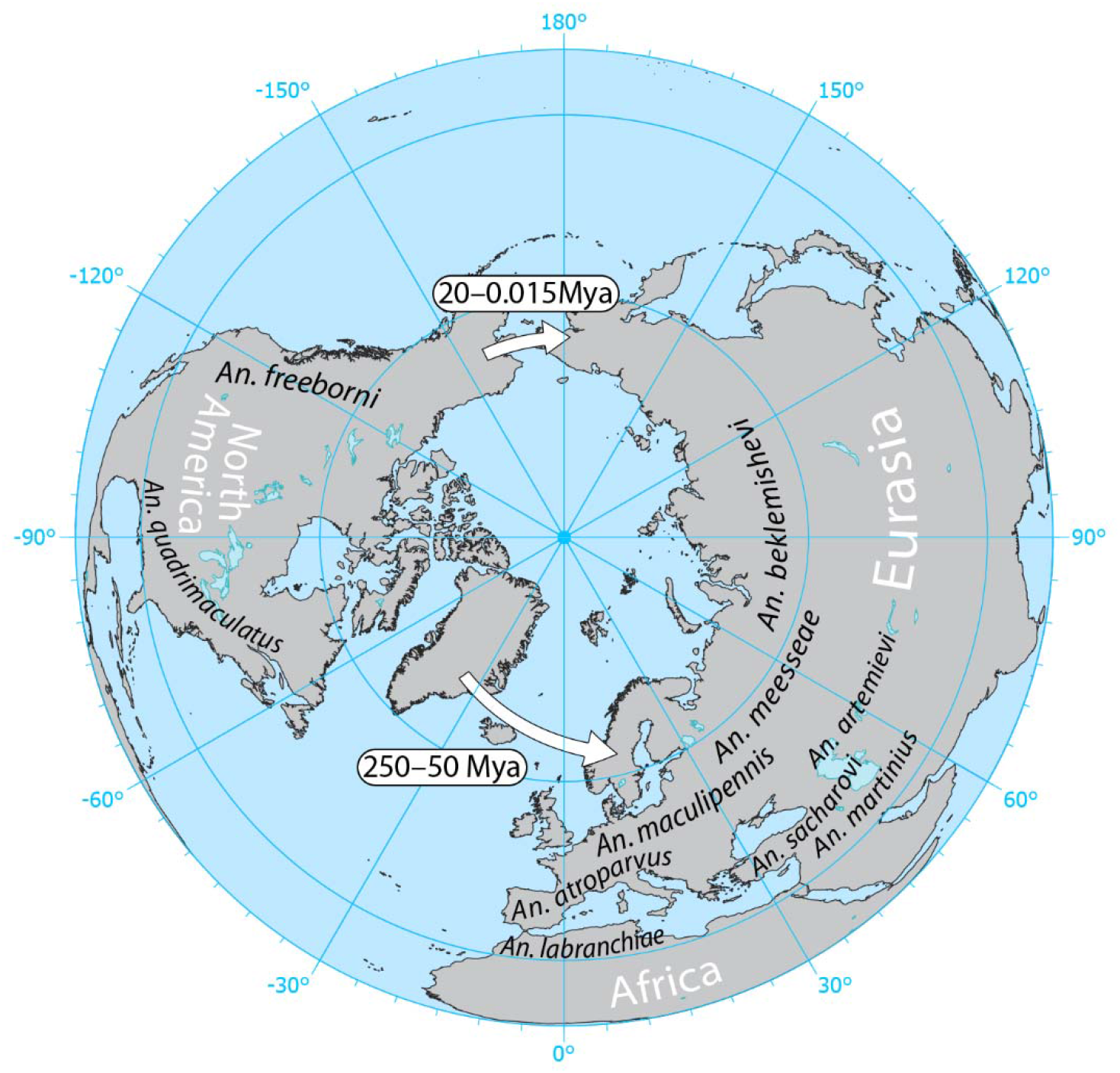
The hypothesis of two migration events of the Maculipennis from North America to Eurasia. The hypothesis was based on phylogenetic relationships between the Maculipennis species revealed by a cytogenetic analysis of their polytene chromosomes. Arrows show possible paths of migration that may have occurred in different times when the two continents were connected through Greenland about 250-50 Mya or through the Bering Land Bridge about 20-0.015 Mya.

The availability of PCR-based molecular technologies allowed researchers to use ribosomal DNA in phylogenetic studies of mosquitoes [32, 33]. Three studies employed Internal Transcribe Spacer 2 (ITS2) sequences of ribosomal DNA to infer phylogenetic relationships between the Palearctic and the Nearctic members of the Maculipennis group [34–36]. These molecular phylogenetic studies provided support for the basal position of species from the Southern Eurasian clade rather than species from the European clade as the cytogenetic studies suggested. Two of the molecular phylogenetic studies included ITS2 sequences from *An. beklemishevi* and pointed to the exceptional position of this species with respect to the Maculipennis subgroup [35, 36] in agreement with the cytogenetic studies. Sequencing of ITS2 in *An. beklemishevi* determined that it is ∼300 bp longer than ITS2 in the other members of the Maculipennis group due to the insertion of a transposable element [35]. Interestingly, the ITS2 sequence-based phylogeny, which included this element in the analysis, grouped *An. beklemishevi* together with the Nearctic species *An. freeborni* and *An. hermsi*. However, if the transposable element was removed from the sequence prior to the analysis, *An. beklemishevi* was placed as the basal branch with the Palearctic members of the Maculipennis group. R. Harbach, in 2004, proposed a classification for the *Anopheles* genus that was largely based on ITS2 sequence analyses [10]. According to this classification, the Maculipennis group was subdivided into two North American subgroups: Freeborni and Quadrimaculatus, and one Eurasian Maculipennis subgroup. *An. beklemishevi*, the species with the most northern distribution in Eurasia, was included in the Quadrimaculatus subgroup distributed in the southeastern USA. However, this placement contradicted the fact that the banding patterns of the polytene chromosomes between this species and *An. earlei* [26, 29] and *An. earlei* belongs to the Freeborni subgroup [10]. More recent molecular studies of ITS2 sequences identified new Palearctic members of the Maculipennis group including *An. artemievi* [37], *An. daciae* [38], and *An. persiensis* [39]. This expanded number of species added to the uncertainties in the phylogeny of the Maculipennis subgroup.

In this study, we address several questions to clarify the historic relationships among the members of the Maculipennis group: (1) Did Palearctic members of the Maculipennis group radiate more recently than the Nearctic species? (2) Which clade, among the European, Northern Eurasian, or Southern Eurasian, is the basal clade within the Maculipennis subgroup? (3) What is the systematic position of *An. beklemishevi*? and (4) What is the likely scenario of species migration and radiation in the Maculipennis subgroup? We used two independent approaches: a genome-wide molecular phylogeny and a rearrangement-based phylogeny centered on the gene orders on the X chromosome and fixed chromosomal inversions in the autosomes. Our study identified phylogenetic relationships within the Maculipennis subgroup, clarifies the historic relationships between North American and Eurasian members, determined the pattern of gene flow between the species, and helped to better understand the route and time of their migration between the continents. The reconstructed phylogeny demonstrates that vectorial capacity and the ability to diapause in the winter evolved multiple times in the evolution of the Maculipennis group.

## Results

### *De novo* genome and transcriptome assemblies of mosquitoes from the Maculipennis group

To generate the phylogeny of the Palearctic members of the Maculipennis group, we sequenced genomes or transcriptomes of 10 Palearctic species and 2 Nearctic species. The genome sequencing reads for 4 species (*An. martinius, An. artemievi, An. melanoon,* and *An. persiensis*) were aligned to the *An. atroparvus* AatrE3 reference genome [40]. The mapping rate was high with 82.08%, 83.4%, 82.11%, and 84.89% of the paired reads properly aligned for *An. martinius, An. artemievi, An. melanoon,* and *An. persiensis*, respectively. We sequenced the transcriptomes of 8 species (*An. messeae, An. daciae, An. quadrimaculatus, An. beklemishevi, An. labranchiae, An. macullipennis, An. freeborni,* and *An. sacharovi*). Transcriptomes from the two geographically distant populations of *An. daciae* in Europe and Asia (Moscow and Tomsk regions, respectively) were also obtained and compared. After transcriptome assembly and annotation using the Transdecoder pipeline, we obtained between 14,404 and 22,326 proteins per species (**Additional file 1: Table S1**), with contig N50 ranging from 681 bp to 1329 bp and the total coding sequence (CDS) lengths ranging from 9.38 to 21.5 Mbp, which are typical transcriptome assembly metrics for non-model organisms. The single-copy orthologs benchmarking against the *Diptera* dataset demonstrated that 28.2 – 73.3% of genes per species were completely assembled during the analysis and 12.8 – 21.8% were fragmented. The duplication level, which can arise as a result of a separate assembly of different haplotypes especially in highly polymorphic species, was generally low and ranged from 0.2% (*An. beklemishevi*) to 3.6% (*An. quadrimaculatus*). Orthofinder analysis returned 1,271 single-copy orthologs, which were present in all 13 species. These genes were aligned and concatenated, resulting in a 1,643,691 bp-long alignment. After the removal of gapped regions, the final alignment consisted of 898,101 bp. JModelTest2 results demonstrated that the best substitution model for our dataset was GTR+G+I, which was used as GTRGAMMAI during RaxML analysis.

### Multigene phylogeny of the Maculipennis subgroup

Maximum-likelihood tree reconstruction using RaxMl demonstrated a high-confidence phylogeny with 100% bootstrap support for all the nodes (**Fig. 2**). The tree was rooted with *An. sinensis* as an outgroup species. The analysis of the tree indicated that Palearctic and Nearctic members of the group represent separate phylogenetic branches. The placement of *An. beklemishevi* together with the Palearctic members of the group argues against the current systematic position of this species within the Nearctic Quadrimaculatus subgroup [10]. According to our phylogeny, *An. beklemishevi* belongs to the Maculipennis subgroup. Moreover, *An. freeborni*, rather than *An. quadrimaculatus*, is more closely related to *An. beklemishevi* and other Eurasian species of the Maculipennis subgroup. After migration of the Maculipennis mosquitoes to Eurasia and separation of the *An. beklemishevi* lineage, they further split into the Southern Eurasian *An. sacharovi*-*An. martinius* clade, ancestors of the European *An. atroparvus*-*An. labranchiae* clade and the Northern Eurasian clade, which includes *An. artemievi*, *An. daciae*, *An. melanoon*, *An. persiensis*, *An. maculipennis*, and *An. messeae*. Interestingly, *An. daciae* from the Moscow region clusters together with *An. daciae* from the Tomsk region rather than with *An. messeae,* which was collected in the Moscow region. This result further supports the species status of *An. daciae* [2].

**Figure 2.**
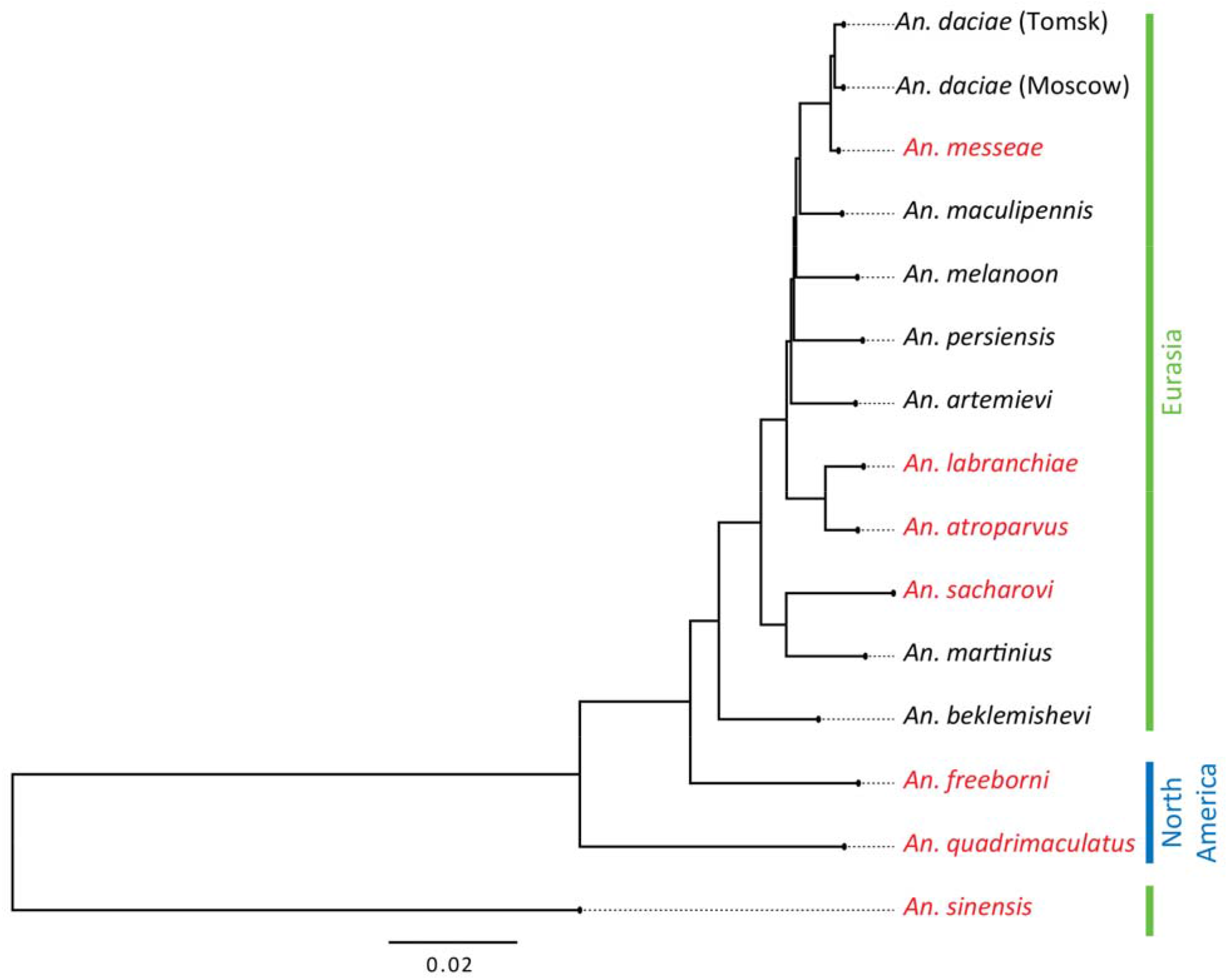
Maximum-likelihood tree of 14 *Anopheles* species based on the 1,271 single-copy orthologs. The major malaria vectors are shown in red. Geographical distribution of the mosquitoes in Eurasia and North America are shown by lines with different colors on the right side of the figure. A scale bar refers to a phylogenetic distance in a fraction of nucleotide differences.

**Figure 3.**
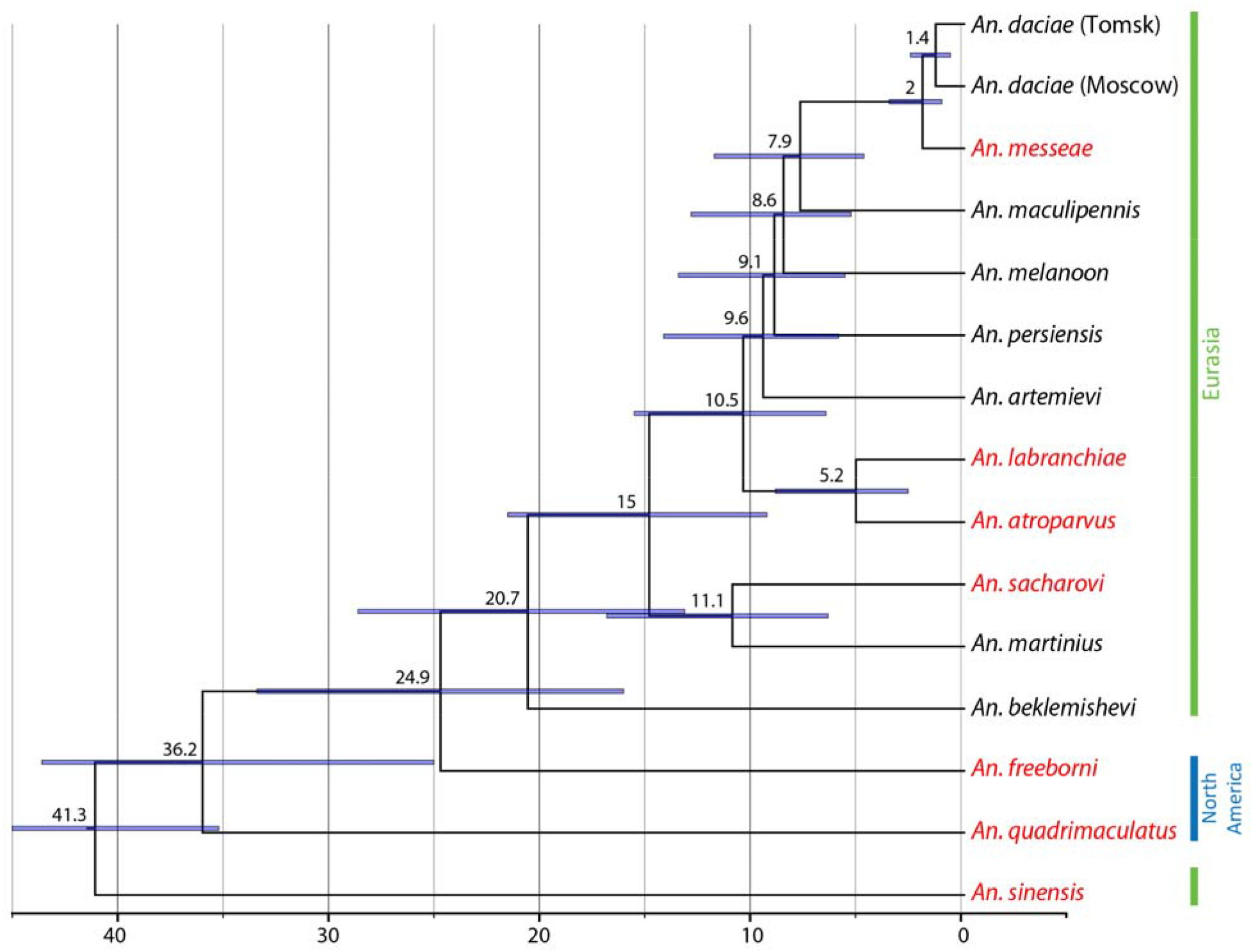
A time-calibrated phylogenetic tree for the studied species based on the 126,025 4-fold degenerated sites. The time scale is in Mya, mean values and time intervals are indicated in blue above the branches. Major malaria vectors are shown in red. Geographical distribution of the mosquitoes in Eurasia and North America are shown by lines with different colors on the right side of the figure.

### Divergence times among species of the Maculipennis subgroup

Our estimation of the divergence times demonstrated that the origin of the Maculipennis group is dated back ∼36.2 Mya when the Quadrimaculatus subgroup separated from the rest of the Maculipennis mosquitoes (**Fig. 2**). The split between the American and Eurasian species of the Maculipennis group occurred between 16 and 34 Mya with a mean estimate of 24.9 Mya. Migration of mosquitoes from North America to Eurasia may have occurred before ∼20.7 Mya in the Neogene Period (Miocene Epoch) followed by a split between the most northern species *An. beklemishevi* and the rest of the species in the Maculipennis subgroup. Around 15 Mya, in the mid-Miocene, a separation between the Southern Eurasian *An. sacharovi-An. martinius* clade and the rest of the species occurred. Next, a split between the European *An. atroparvus-An. labranchiae* clade and the Northern Eurasian clade took place around 10.5 Mya. The most recent species divergence occurred between *An. daciae* and *An. messeae* between 3.6 and 1.1 Mya (a mean estimate of 2 Mya) in the Pliocene-Pleistocene Epoch at the end of the Eogene Period. The mean divergence time of 1.4 Mya between the Tomsk and Moscow populations of *An. daciae* is probably an overestimation as population divergence time can be better assessed using population samples and coalescent analysis, taking into account migration events and demographic features of populations [41].

### Signatures of genomic introgression between the species of the Maculipennis group

To understand the pattern of a possible gene flow and introgression events, implying absence of complete reproductive isolation among members of the Maculipennis group, we used the D statistics (ABBA-BABA test, 4-taxon) analysis (**Fig 4**). Our study detected highly significant and widespread signatures of introgression events between the species within the group including species which are geographically isolated today such as *An. labranchiae* and *An. beklemishevi* or *An. freeborni* and *An. sacharovi*. The current phylogenetic setting of the analysis cannot unfortunately assess the level of introgression between closely related species such as *An. labranchiae* and *An. atroparvus* where population sampling is needed.

**Figure 4.**
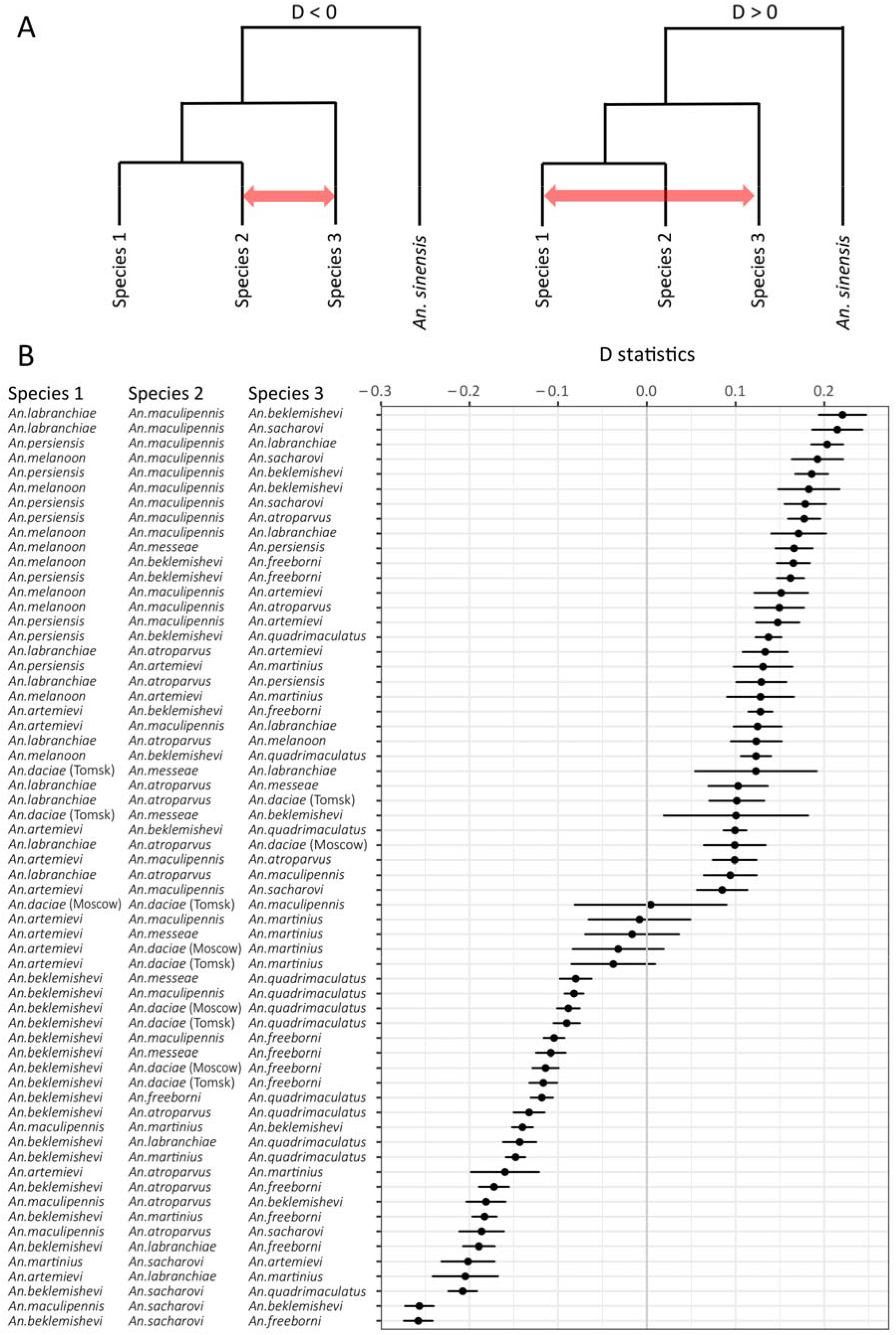
Signatures of genomic introgression among the species of the Maculipennis group. A) A schematic explanation of the analysis using D statistics. We tested for discordant genealogies given the maximum-likelihood species tree between species 2 and 3 and species 1 and 3. B) Introgression statistics (D) among species shown on left and their standard deviations (SD) shown by lines on right. Only combinations of species with Bonferroni corrected P-values < 0.001 are shown. Negative values of D statistics imply gene flow between species 2 and 3, positive values between species 1 and 3, with *An. sinensis* as an outgroup as schematically depicted in panel A.

### Chromosome phylogeny of the Maculipennis subgroup

To understand the karyotypic evolution within the Maculipennis subgroup, we analyzed X chromosome rearrangements among five members and autosomal rearrangements among eight members. The banding patterns of X chromosomes could not be compared visually due to the high numbers of fixed rearrangements among the species [26, 28]. For this reason, the X chromosomal rearrangements were identified based on the order the genes mapped using fluorescence *in situ* hybridization (FISH). We selected 21 genes from the *An. atroparvus* genome [40] located from 208,734 bp to 1,320,538 bp (∼800 kb on average) according to the genome map (**Additional file 1: Table S2**). The DNA probes were designed based on the exons of these genes.

The probes were amplified, labeled, and hybridized to the polytene chromosome preparations from ovarian nurse cells of *An. atroparvus* to confirm the order and distances between them on the X chromosome (**Fig. 5A**). Selected markers benchmarked ∼75% of the physical length of the X chromosome from subdivisions 1A to 4B, representing the euchromatic part of the chromosome [42]. We hybridized the same probes with polytene chromosomes of *An. beklemishevi, An. labranchiae*, *An. maculipennis*, and *An. sacharovi*. Not all of the genes were successfully mapped to the chromosomes of all the species. We were able to hybridize and map all 21 genes only in *An. labranchiae* and *An. maculipennis,* 20 and 19 DNA probes hybridized to the chromosomes of *An. sacharovi* and *An. beklemishevi*, respectively. In total, 17 gene probes hybridized to all the analyzed species and were used for further analysis. Gene orders in three species, *An. labranchiae*, *An. atroparvus*, and *An. maculipennis,* were identical, while gene orders in the remaining species were different from each other. Differences among species in the chromosomal position of the gene marker AATE17741 are shown in **Fig. 5B**.

**Figure 5.**
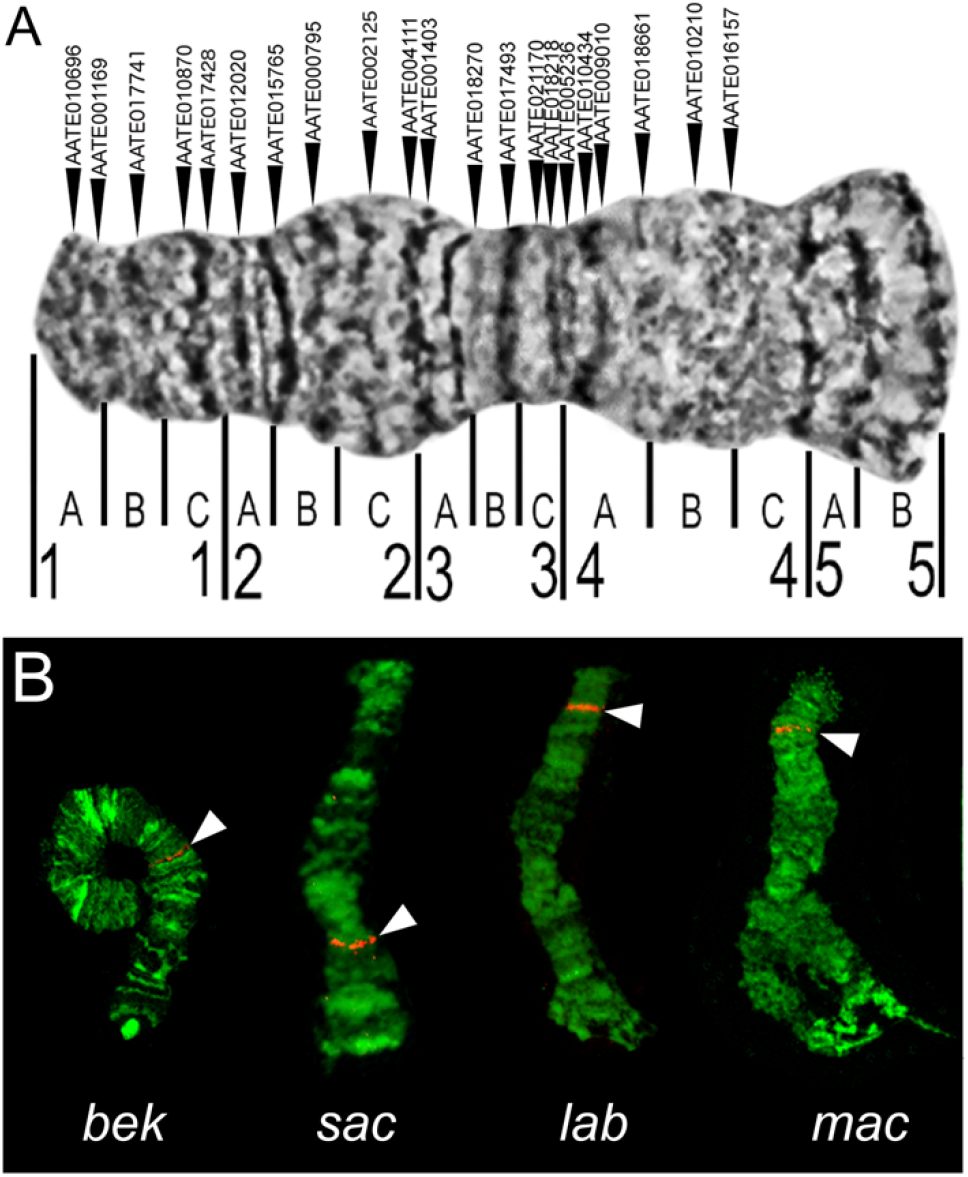
Physical mapping of marker genes on X chromosomes of species from the Maculipennis group. Positions of 21 marker genes in the X chromosome of *An. atroparvus* (A) are shown by arrows above the chromosome. Numbers and letters below the chromosome indicate numbered divisions and lettered subdivisions. Localization of the gene AATE017741 in the Х chromosome of *An. beklemishevi* (*bek*), *An. sacharovi* (*sac*), *An. labranchiae* (*lab*), and *An. maculipennis* (*mac*) are shown in panel B, indicating significant reshuffling of chromosome arrangements among the species.

We calculated the number of X chromosome rearrangements and reconstructed phylogenetic relationships among seven species using the Multiple Genome Rearrangements (MGR) program [43]. Conserved gene orders were considered as synteny blocks. The pattern of synteny blocks in *An. atroparvus* was considered as the standard. The following orders and orientations of conserved genes and synteny blocks were used as an input for the MGR analysis.

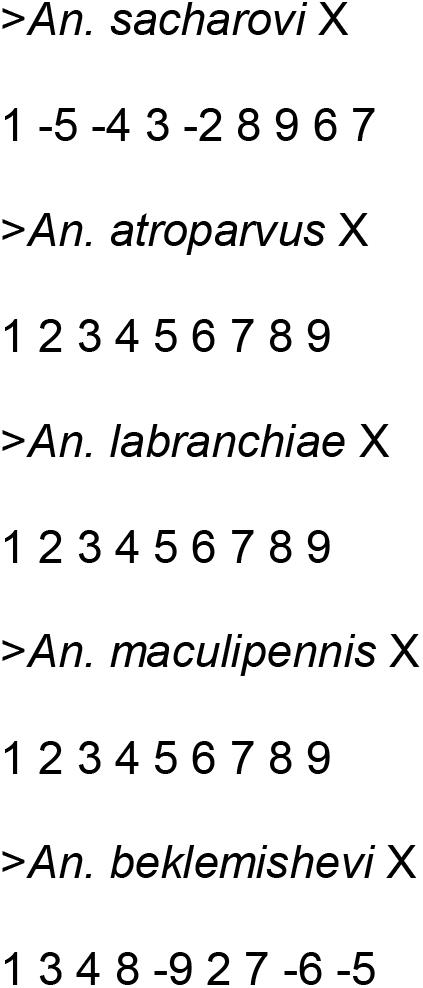

The MGR program reconstructed putative ancestral karyotypes and created a phylogenetic tree by implementing an algorithm that minimized the sum of the rearrangements over all edges of the phylogenetic tree (**Fig. 6A**). The program clustered together *An. atroparvus*, *An. labranchiae,* and *An. maculipennis* since they had no fixed inversions. *An. sacharovi* was separated from *An. atroparvus* by four fixed inversions. Overall, the X chromosome rearrangement topology agrees with the whole-genome molecular phylogeny demonstrating that *An. beklemishevi* is the most distantly related species to the other members.

**Figure 6.**
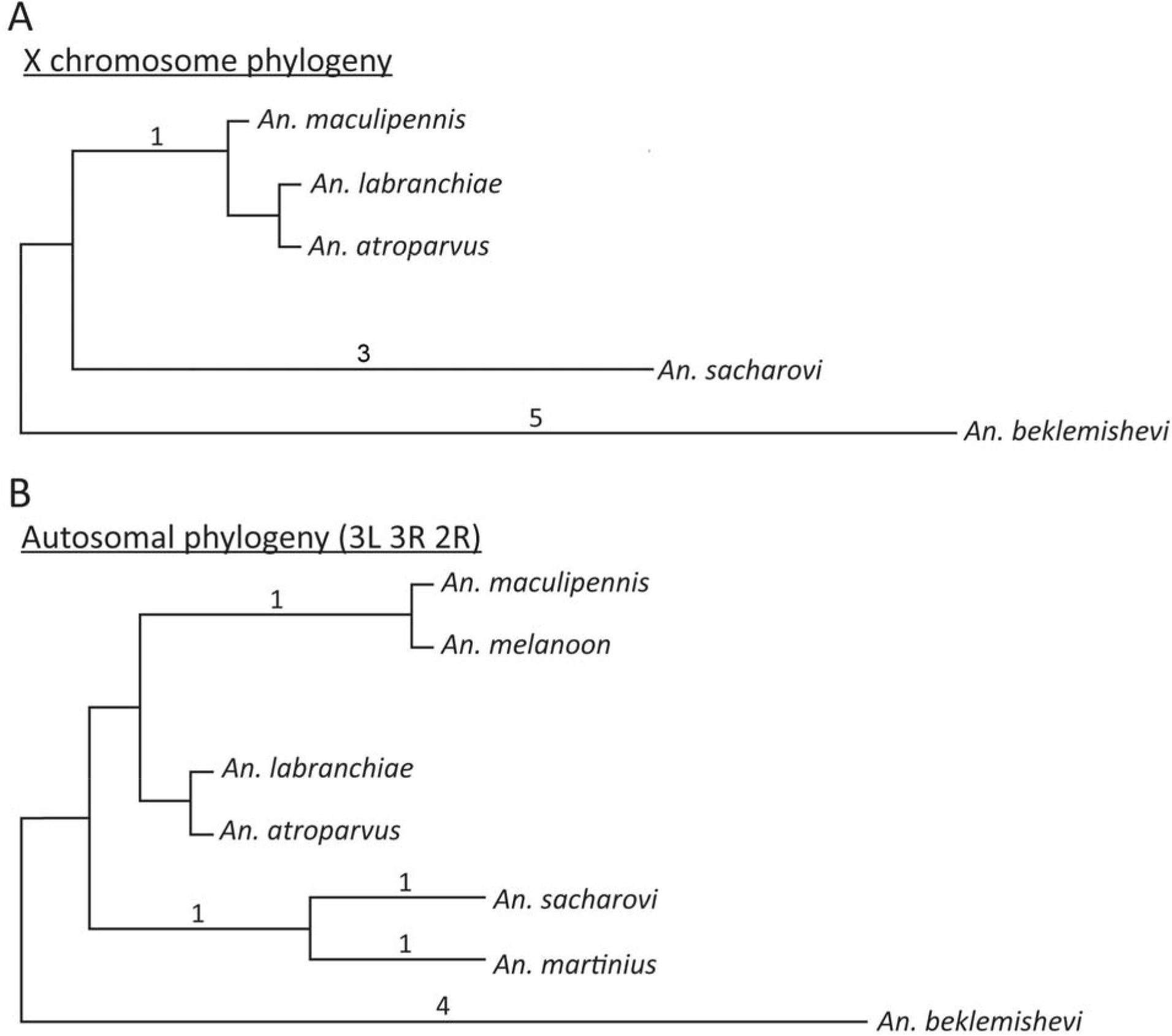
Chromosome-based phylogeny for the Maculipennis group. (A) Rearrangement phylogeny based on positions of 17 DNA probes mapped to the X chromosomes of five species. (B) Rearrangement phylogeny based on autosomal banding pattern differences detected in eight species. Numbers above tree branches indicate the number of inversions fixed between the lineages.

We also analyzed rearrangements on the autosomes in eight Palearctic members of the Maculipennis subgroup. For this analysis, we used the chromosome banding patterns that were previously described by V.N. Stegniy [21, 26]. Chromosomes of *An. atroparvus*, *An. beklemishevi, An. labranchiae*, *An. maculipennis*, *An. martinius*, *An. melanoon*, *An. messeae*, and *An. sacharovi* were compared. Banding patterns in *An. atroparvus* were considered as the standard. Chromosomal rearrangements were determined in the 2R, 3R, and 3L autosomal arms. Banding patterns in two groups of species were identical: *An. atroparvus*-*An. labranchiae* and *An. maculipennis*-*An. melanoon-An. messeae*. The following orders and orientations of conserved synteny blocks on autosomes were used as an input for the MGR analysis.

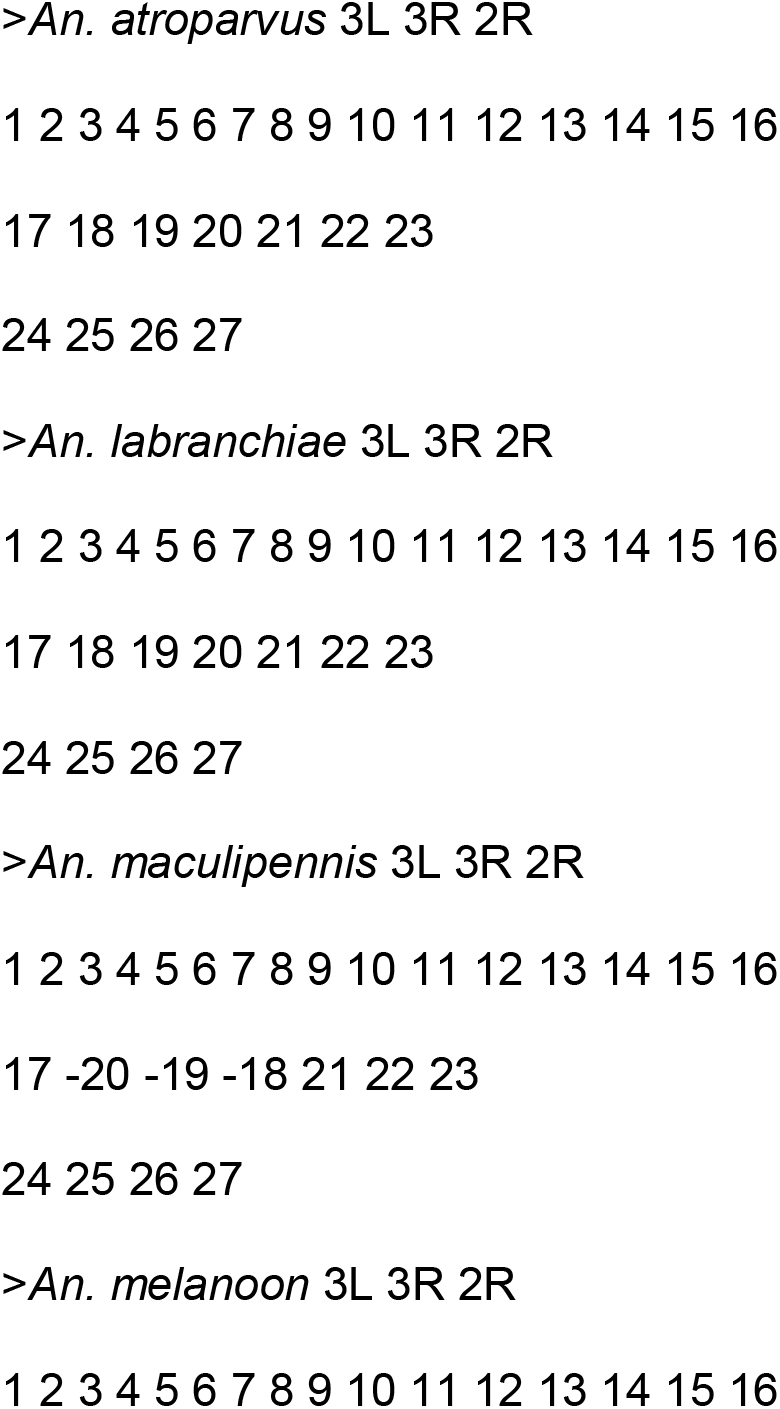
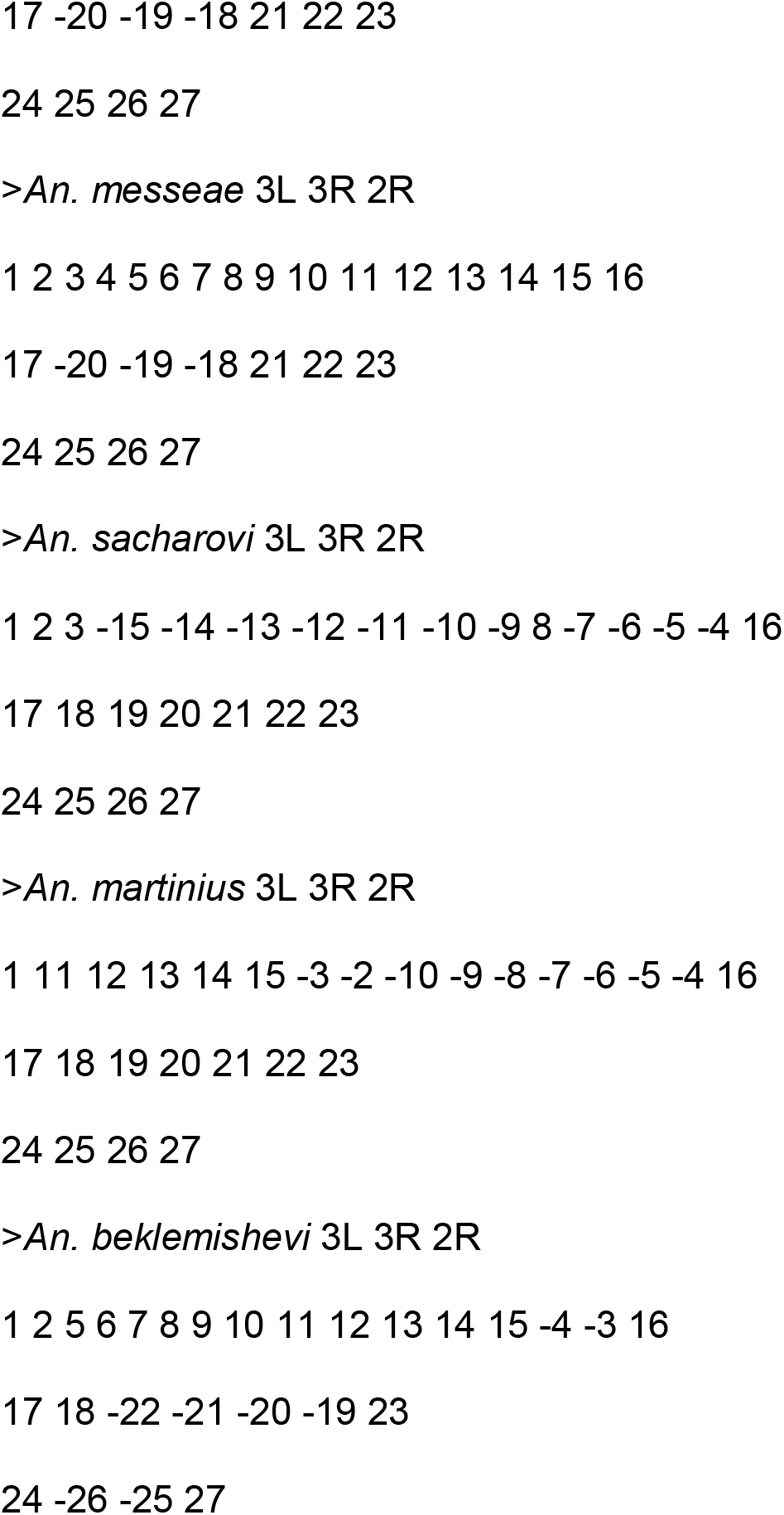

Overall, the topology of autosomal (**Fig. 6B**). and X chromosome phylogenetic trees (**Fig. 6A**) were very similar to each other. *An. atroparvus* and *An. labranchiae* clustered together. The *An. maculipennis-An. melanoon-An. messeae* cluster was separated from the *An. atroparvus*-*An. labranchiae* cluster by one inversion. *An. sacharovi* and *An. martinius* were separated from *An. atroparvus* by two inversions. In agreement with the X rearrangement and the molecular phylogenies, *An. beklemishevi* was the most distant from the rest of the species.

## Discussion

### The phylogeny of the Maculipennis subgroup

In this study, we reconstructed a multigene phylogeny for the Palearctic members of the Maculipennis group. Our phylogeny, for the first time, included all 11 described Eurasian species together with two North American species and demonstrated that the Palearctic members of the Maculipennis group radiated more recently. Similar to the chromosomal [26, 28] and ITS2-based phylogenies [35], our analysis supported the presence of three major clades in the Maculipennis subgroup: the Southern Eurasian clade of *An. sacharovi* and *An. martinius,* the European clade of *An. atroparvus* and *An. labranchiae*, and the Northern Eurasian clade of the remaining species. Similar to the chromosomal rearrangement phylogeny [26, 28] and the ITS2-based phylogenies [34–36], our phylogeny clusters together species with identical chromosomal patterns, *An. atroparvus* and *An. labranchiae* [17]. Unlike the ITS2-based phylogenies [34–36], our genome-wide phylogeny provides strong support for the branching order of all Eurasian species. The Southern Eurasian clade represents a basal phylogenetic branch with respect to the European clade and the Northern Eurasian clade. Our autosomal and X chromosomal rearrangement phylogenies were similar to the multigene phylogeny and supported the most distant position of *An. beklemishevi* with respect to the other members of the Maculipennis subgroup.

The systematic position of *An. beklemishevi* remained unclear until our study. The current taxonomic nomenclature placed *An. beklemishevi*, a species with a broad distribution in the most northern areas of Eurasia, into the Nearctic Quadrimaculatus subgroup [10]. This subgroup includes the dominant malaria vector *An. quadrimaculatus* [14] and four other species. *An. quadrimaculatus* has a broad distribution in the eastern and midwestern areas of the USA. Distribution of other species from the Quadrimaculatus subgroup is restricted to Florida. The Nearctic Freeborni subgroup includes *An. freeborni,* a dominant vector of malaria in the Western United States [14] and three other species: *An. occidentalis, An. hermsi,* and *An. earlei* [10]. Distribution of *An. freeborni*, *An. occidentalis*, and *An. hermsi* is restricted to the west coast of the North America. In contrast, *An. earlei* has a broad distribution in the northern areas in North America. The cytogenetic analysis determined differences in chromosomal banding patterns caused by the overlapping chromosomal inversions between the Palearctic members of the *An. maculipennis* group [17–19]. The phylogenetic position of *An. beklemishevi* remained unresolved because of the absence of an intermediate chromosome arrangement that could connect *An. beklemishevi* with other species. Nevertheless, comparison of its chromosomes with Nearctic members of the group suggested that *An. beklemishevi* is a close relative to *An. earlei* [29]. Our genome-based and rearrangement-based phylogenies strongly suggest that *An. beklemishevi* belongs to the most basal branch among the Palearctic members in the Maculipennis group. Moreover, the Eurasian species are more closely related to *An. freeborni* than to *An. quadrimaculatus*. Therefore, our study argues against placing *An. beklemishevi* into the Quadrimaculatus subgroup. Phylogenomic analysis of the remaining North American species will help to better resolve the systematic positions of all members of the Maculipennis group.

Our phylogenetic tree allowed for the analysis of introgression events between the species. The introgression events were detected between many members of the group. For example, *An. beklemishevi* demonstrated significant signatures of introgression with allopatric species *An. labranchiae.* Also, the Nearctic species *An. freeborni* showed significant signatures of introgression with the Palearctic species *An. sacharovi*. The evidence of gene flow between allopatric species can be interpreted as historical events occurred during the last Glacial Maximum in refugia regions when the species boundaries were shifted and differed significantly from the current distribution. Another explanation can be the current gene flow events across the species boundaries which permit genetic exchange between the chain of species and their populations. In this situation two allopatric species can demonstrate genetic admixture indirectly mediated by a third species which is sympatric for both of them. To prove the existence of such a scheme one will need to sample major populations from diverse geographic regions of several species. The phenomena of widespread historical introgression and current incomplete reproductive isolation has been demonstrated in the African *An. gambiae* complex [44] using whole genome datasets. It is possible that genetic introgression can be a factor contributing to the acquisition of adaptations related to malaria vectorial capacity. For example, a study of introgression between members of the *An. gambiae* complex suggested that traits enhancing vectorial capacity can be acquired from nonsister vector species through a rapid process of interspecific genetic exchange [44]. Also, a genomic analysis of the *An. funestus* complex demonstrated substantial gene flow into the malaria vector *An. funestus* before this species expanded its range across tropical Africa [45].

Among 13 studied members of the Maculipennis group, 6 species *An. messeae*, *An. labranchiae*, *An. atroparvus*, *An. sacharovi, An. freeborni,* and *An. quadrimaculatus* are dominant vectors of malaria [13, 14]. However, the malaria vectors do not form a monophyletic group. Thus, our phylogenomic analysis demonstrated that vectorial capacity evolved multiple times in the evolution of the Maculipennis group and likely depended on species distribution rather than on phylogenetic relationships.

### A working hypothesis of species radiation, migration, and adaptation in the Maculipennis subgroup

Based on the current distribution of species from the Maculipennis group and the phylogenies developed in this study, we propose that migration of the Maculipennis mosquitoes occurred through the Bering Land Bridge ∼20.7 Mya during the Miocene. Because of the fluctuation of the sea level, this bridge existed intermittently across the Bering Straits, connecting Chukotka with Alaska and allowing animals to migrate between North America and Eurasia [30]. The connection between the two continents occurred from the Paleocene, ∼60 Mya, through the Eocene, ∼40 Mya, and from the Miocene, ∼20 Mya, until relatively recent times. Various species of animals [46] including horses [47] and most recently ancient humans [48, 49] were able to use the Bering Land Bridge for their migration between the continents. The possible alternative scenario of the Maculipennis group migration through Greenland is unlikely. It was originally proposed to explain the distribution of the majority of the species in Europe and Western Asia rather than in Eastern Asia and the Far East. Although, the connection between North America and Eurasia through Greenland existed in the Paleocene ∼60 Mya, it was disrupted in the Eocene [30] before mosquitoes migrated according to our phylogeny. Moreover, the basal species among Eurasian Maculipennis mosquitoes, *An. beklemishevi*, is more closely related to *An. freeborni,* the species from the western part of North America located close to the Beringia, rather than to eastern in North American species *An. quadrimaculatus*.

We think that evolution of the Maculipennis subgroup have been affected by multiple glaciation events occurred in Eurasia in the Neogene Period after 23 Mya [30]. We speculate that glaciation shaped the divergence and geographic distribution in the Palearctic Maculipennis species by pushing their natural ranges westward (**Fig. 7**). At the last glaciation maximum, the ice sheet covered northern Europe, Scandinavia, northeastern Asia, and a large part of North America [50]. Moreover, much of Siberia, the Far East, and Central Asia were desert-like with only ∼2% of the ground covered by vegetation. The ice sheet distribution during the last glaciation period may explain the current absence of Maculipennis mosquitoes from the Eastern territories of Eurasia. We hypothesize that the ancestral species of the Maculipennis subgroup arrived in Eurasia with a well-developed ability to diapause during winter. This species may have given rise to *An. beklemishevi*, which remained in the northern territories but had to move westward. The origin of the Southern Eurasian clade, *An. sacharovi*-*An. martinius,* occurred when the overall temperature increased in the middle of the Miocene, 15 Mya [30]. These species lost their ability to diapause and developed the ability to survive in hot and arid climates. Other species reached Europe ∼10.5 Mya and gave rise to the European *An. atroparvus*-*An. labranchiae* clade and the Northern Eurasian clade. Among the species of the Northern Eurasian clade, *An. artemievi*, *An. persiensis,* and *An. melanoon,* adapted to the warmer climate. Around 8 Mya, when the planet temperature was cooling again, *An. maculipennis, An. messeae,* and *An. daciae* of the Northern Eurasian clade developed an obligatory diapause and became sympatric with *An. beklemishevi*. All the species from this clade gradually moved toward the east. Interestingly, in species with a large distribution in Eurasia, such as *An. maculipennis,* northern populations can enter a deep diapause to overwinter but southern *An. maculipennis*, like the southern species *An. sacharovi*, preserves its ability to bloodfeed during winter [51]. The most recent split between *An. messeae* and *An. daciae* overlapped with the strong glaciation period that occurred ∼3 Mya [30]. Because of currently rising temperatures, the southern species of the Maculipennis subgroup have been spreading northwards. For example, *An. maculipennis* recently extended its range to the northeast and has reached the Southern Urals [52]. Overall, these observations suggest, that obtaining or losing the ability to diapause in winter may evolved multiple times in the evolution of the Maculipennis group depending on the climate conditions.

**Figure 7.**
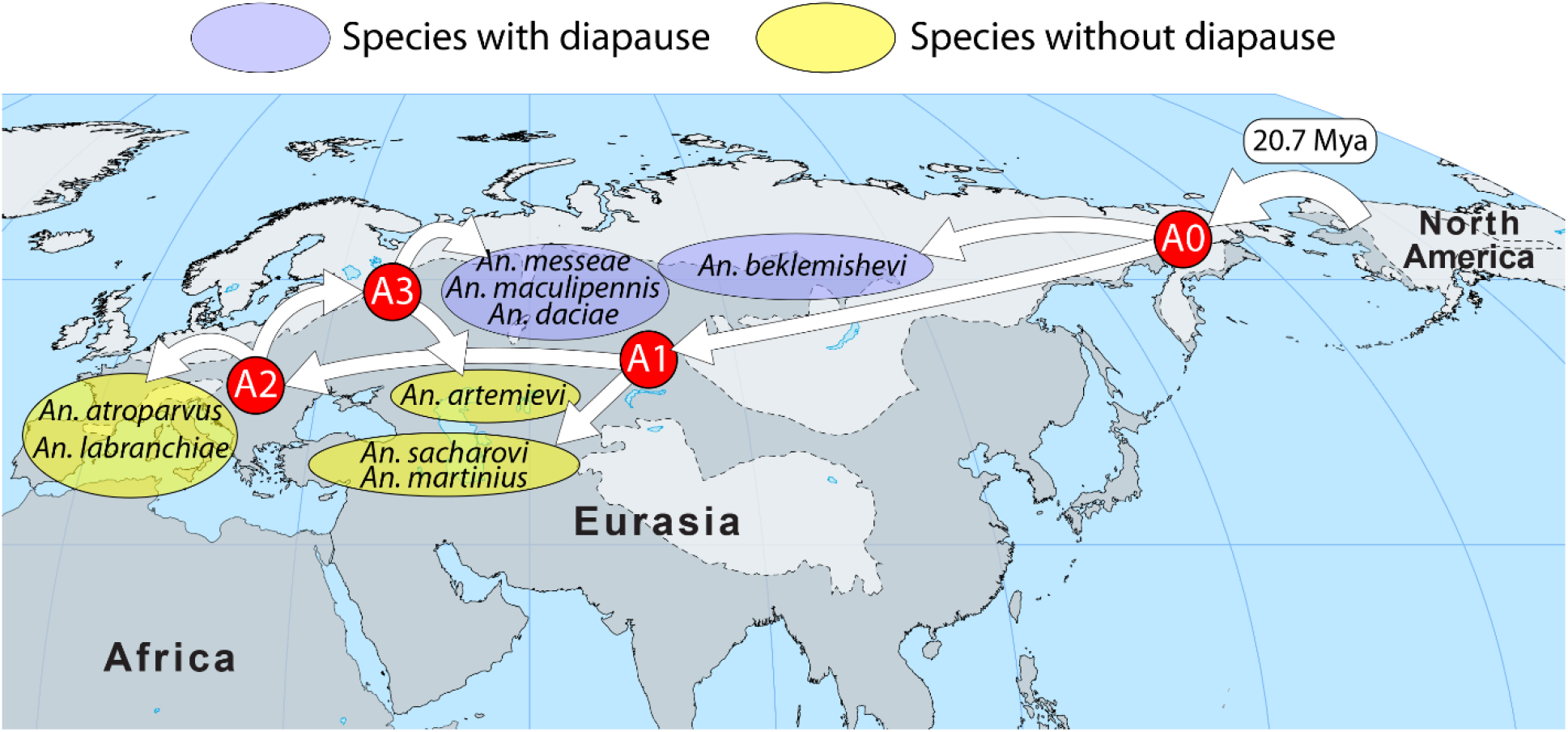
A scenario of the Maculipennis group migration through the Bering Land Bridge. This scenario is supported by existence of the connection between Alaska and Chukotka at the mosquito migration time ∼20.7 Mya. The last large glaciation (shown in light grey color) explains why the Maculipennis mosquitoes are not present in the Far East. Letters A0, A1, A2, and A3 in red circles represent hypothetical ancestral species for the phylogenetic clades. Diapausing and non-diapausing species are shown in blue and yellow ovals, respectively.

Our study suggests that adaptation of malaria mosquitoes to different climates was an important factor in their evolution. Radiation of the Maculipennis subgroup in Eurasia coincided with the highest rate of overall speciation among Anophlinae mosquitoes in the world, which occurred around 20 Mya [4]. The increase of speciation in mosquitoes strongly correlates with the overall increase in concentration of atmospheric CO_2_ and in overall temperature on the planet [53]. The elevated mosquito diversification also overlapped with the evolution of grasses [54], which provided new habitats for the larval stages of mosquitoes and facilitated speciation of mammal species, such as ruminants [55] and rodents [56], which increased the host availability for adult mosquitoes [53]. The current change in climate may lead to geographical redistribution of malaria vectors and return malaria transmission back to territories where it was eradicated [57].

## Conclusions

In this study, we reconstructed the relationships among the Eurasian members of the Maculipennis group using multigene and chromosome phylogenies. The analysis has demonstrated that genome-wide approaches are highly effective for reconstruction of the evolutionary history of the Holarctic malaria mosquitoes. Our genome-wide phylogeny provided strong support for the branching order of all Eurasian species and for a basal position of the Southern Eurasian clade with respect to the European and Northern Eurasian clades. We demonstrated that *An. beklemishevi* clusters together with the other Eurasian members and, thus, belongs to the Palearctic Maculipennis subgroup. Phylogenomic data support migration of the Maculipennis mosquitoes from North America to Eurasia through the Bering Land Bridge ∼20-25 million years ago. Finally, our study indicated that vectorial capacity and the ability to diapause evolved multiple times in the evolution of the Maculipennis group.

## Materials and Methods

### Mosquito species studied, sample collections, and species identification

The aim of our study was to reconstruct the whole-genome phylogeny of the Maculipennis subgroup. For this purpose, we included 14 species of malaria mosquitoes: two North American species (*An. freeborni* and *An. quadrimaculatus*) and 12 Eurasian species (*An. artemievi, An. atroparvus*, *An. beklemishevi*, *An. daciae*, *An. labranchiae*, *An. maculipennis, An. martinius, An melanoon, An. messeae*, *An. persiensis*, *An. sacharovi*, and *An. Sinensis* (Wiedemann, 1828), an outgroup species). For genome and transcriptome sequencing, we sampled 12 species (*An. artemievi, An. beklemishevi*, *An. daciae* (Moscow and Tomsk populations), *An. labranchiae*, *An. maculipennis, An. martinius, An melanoon, An. messeae*, *An. persiensis*, and *An. sacharovi*) and two North American species (*An. freeborni* and *An. quadrimaculatus*) (**Additional file 1: Table S3**). The genome sequences of *An. atroparvus* EBRO (AatrE3) and *An. sinensis* (AsinS2) were retrieved from VectorBase [58]. Sanger sequencing of ITS2 was performed on the collected mosquitoes to identify species. PCR reactions and primer choice for ITS2 sequencing were conducted in accordance with the previous study [59]. PCR products were purified using the Wizard PCR Cleanup and Gel Purification system (Promega, Fitchburg, WI, USA).

### Transcriptome and genome sequencing

We performed transcriptome sequencing for *An. messeae, An. daciae* (Moscow and Tomsk populations)*, An. quadrimaculatus, An. beklemishevi, An. labranchiae, An. macullipennis, An. freeborni,* and *An. sacharovi*. Blood-free females and males of *Anopheles* mosquitoes were fixed in RNAlater to prevent RNA degradation. Total RNA was isolated from 10 individual specimens from each of the 8 species. The polyA-selection method was used and quality control of the RNA sequencing libraries was performed, including size evaluation by a bioanalyzer and by quantitative assay (Fasteris, Inc., Geneva Swiss). Transcriptome sequencing was done using HiSeq 2500 1×100 or 1×125 bp. We received from 8.2 to 12.56 Gb of data for each sample. For the genome sequencing of *An. martinius, An. artemievi, An. melanoon,* and *An. persiensis*, DNA was extracted from adult or larvae mosquitoes using the DNeasy Blood and Tissue extraction system (Qiagen, Germantown, MD, USA) following the manufacturer’s protocol with slight modifications. Genomic pools were created for each species and then sequenced using the Illumina HiSeq 4000 Platform PE150. We obtained from 10× to 70× genome coverage for each species.

### Transcriptome assembly and annotation

The raw reads were trimmed and the remains of adapters were removed using Trimmomatic software [60] with the following settings: *ILLUMINACLIP:2:30:10 LEADING:24 TRAILING:24 SLIDINGWINDOW:4:24 MINLEN:50*. The transcriptomes were assembled using Trinity 2.1.1 software [61] with strand-specific settings. Open reading frames (ORFs) were extracted from the assemblies with Transdecoder utility [62] and translated ORFs were blasted against a collection of mosquito proteins from VectorBase [58] with an e-value threshold 1e-5 using the blastp software [63]. Proteins with blast hits were used in the TransDecoder. Then, the Predict module was employed to find additional coding sequences that did not have any blast hits within datasets based on Hidden Markov Model (HMM) profiles derived from the training set of proteins homologous to other mosquitoes. To remove redundancy caused by the possible assembly of different isoforms, haplotypes, and fragments of genes, the resulting sets of proteins were clustered using CD-HIT software [64] with a 97% identity threshold *(-c 0.97 -n 5 – M 2000*). Thus, the final sets of proteins consisted of collapsed proteins with either blast hits against the *Anopheles* proteins or predicted using TransDecoder, Predict, or the HMM-based module algorithm. The corresponding mRNA and CDSs were extracted from each species and subsequently used for the downstream analysis. To assess the quality and completeness of the assembled transcriptomes, benchmarking of single-copy orthologues using BUSCO V.1.22 [65] was applied with a Diptera-specific dataset consisting of 2,799 single-copy genes and the following settings: *--m OGS --species aedes*.

### Pseudoassembly of whole-genomic sequences

Genomic reads were aligned to the *An. atroparvus* AatrE3 genome using Burrows-Wheeler Aligner (BWA) mem software [66]. The resulting reads were sorted using Samtools [67]. Binary Alignment Map (BAM) files were used to create consensus sequences with the following settings: *samtools mpileup -q 50 -Q 26 | bcftools call | vcfutils.pl vcf2fq -d 8 -D 80*. To reduce possible errors caused by interspecies alignment, stringent criteria were used during the consensus construction including a minimum phred-scaled mapping quality equal to 50 (*-q* 50), a minimal phred-scaled quality of a base to call a substitution equal to 26 (*-Q* 26), and minimum and maximum depths equal to 8 and 80, respectively. The maximum depth threshold was utilized to reduce the likelihood of calling substitutions in highly duplicated or repetitive regions in addition to the mapping quality threshold. During assembly, genomic regions that were not covered or poorly covered were converted into Ns and excluded from subsequent analysis. The consensus transcripts were processed with seqtk utility (https://github.com/lh3/seqtk) to select a random base from heterozygous positions and then translated into amino acid sequences for the orthology inference step.

### Orthology inference and alignment

Amino acid sequences were assigned to the orthogroups using OrthoFinder 1.1.4 software [68] using the total set of 15 species. Orthogroups were manually filtered to exclude orthogroups containing missed species and any paralogs. The nucleotide sequences from the single-copy orthogroups (CDSs) were aligned in the codon-aware mode (*F+ -codon*) using Prank software [69]. The aligned sequences were concatenated and cleaned using Trimal [70] utility, which removed all codons with any gaps in alignment (*-nogaps*).

### Phylogenetic tree reconstruction

To determine the optimal substitution model for the dataset, jmodeltest2 [71] software and Bayesian Information Criteria (BIC) were utilized. The maximum likelihood tree was constructed using the RaxML 8 package [72] and the GTRGAMMAI model. The topology was tested with 1000 bootstrap replications. To determine the divergence time of the main Maculipennis lineages, we extracted 126,025 4-fold degenerated sites using MEGA 7 software [73] from the final alignment. The divergence time was estimated using MCMCtree from the PAML 4.8 [74] package using the approximate likelihood calculation and the GTR+G model of nucleotide substitution after 10 million MCMC generations (including the first one million being discarded as a burn-in). Due to the lack of well-established fossil calibrations for the studied and closely related taxonomic groups, we used a root calibration reflected divergence of *An. sinensis* for the rest of the group (from 35 to 45 million years ago) based on previous work [9].

### Analysis of introgression

To test our hypothesis for interspecies introgression, we used Hybridcheck software [75]. This software uses well-established statistics: D statistics (ABBA-BABA test) [76] and the f statistic [77] to test hypotheses about phylogenetic discordance caused by putative hybridization events for 4-taxon sets. The significance of the D statistic was tested using the jackknife method with 10 blocks and then corrected using the Bonferroni approach. The software generates all possible 4 taxon trees and calculates statistics for each of them. After the calculation, all significant results were manually checked for correspondence to the maximum-likelihood phylogeny and discordant trees and redundant species combinations were removed from the analysis.

### Chromosome preparation

Polytene chromosome preparations for fluorescence *in situ* hybridization (FISH) were prepared from ovarian nurse cells of female malaria mosquitoes from the following species: *An. atroparvus*, *An. beklemishevi*, *An. labranchiae*, *An. maculipennis*, and *An. sacharovi*. Adult mosquito ovaries were dissected and fixed in Carnoy’s solution (3 parts 96% ethanol to 1 part glacial acetic acid). Ovaries were stored at -20C° for up to one year before using. For squashed chromosome preparations, 20-30 follicles were put on a glass slide in a drop of ice-cold propionic acid and left for 5 minutes for maceration. The drop of propionic acid was then changed by a new one, follicles were spread along the slide using needles, covered with an 18x18 coverslip, and squashed by tapping the coverslip using the needle handle. Preparations were then covered by a piece of filter paper for additional tapping. The quality of the chromosome spreading was analyzed under phase contrast microscopy using AxioImager A1 (Carl Zeiss, OPTEC, Novosibirsk, Russia) and AxioVision 4.8.1. software (Carl Zeiss, OPTEC, Novosibirsk, Russia). Only high-quality preparations were used for further procedures. Coverslips were removed by a razor blade after freezing the preparation by dipping it in liquid nitrogen and then slides were immediately placed in ice-cold 50% ethanol for 5 minutes. Subsequently, slides were kept in 70% and 96% ethanol for 5 min and air-dried. After one-week storage at room temperature, chromosomal preparations were used for FISH.

### DNA probe preparation and fluorescence *in situ* hybridization

Exons of 21 selected genes from the *An. atroparvus* genome were used to design primers using the Primer-BLAST tool [78]. Sequences of expected PCR products were matched against the *An. atroparvus* genome using the VectorBase BLAST tool to ensure uniqueness of the DNA-probes. DNA-probes were amplified by PCR and labeled using the Random Primer Labeling Protocol, as described earlier [79]. The DNA-probes were precipitated in ethanol, dissolved in hybridization solution (50% formamide, 10% sodium dextran sulfate, 0.1% Tween-20 in 2× Saline-Sodium Citrate (SSC), pH 7.4) and stored at -20 until use. FISH was performed following our standard protocol [80, 81]. Microscopic analysis of chromosome preparations after FISH was performed using AxioImager Z1 (Carl Zeiss, OPTEC, Novosibirsk, Russia) equipped with a device for improving images, ApoTome (Carl Zeiss, OPTEC, Novosibirsk, Russia) and a CCD camera MRm (Carl Zeiss, OPTEC, Novosibirsk, Russia). Images were captured and processed using AxioVision 4.8.1. software (Carl Zeiss, OPTEC, Novosibirsk, Russia).

### Multiple Genome Rearrangement analysis

The calculation of inversion distances among the included species of the Maculipennis group species was performed using the Multiple Genome Rearrangement (MGR) program available at http://grimm.ucsd.edu/MGR/. The signed option of the MGR program was used. This program implements an algorithm that uses a parsimony approach, *i.e*. it minimizes the sum of the rearrangements over all the edges of the phylogenetic tree [43]. To create an inversion phylogenetic tree, numbers were assigned to represent each conserved synteny block in species using our gene mapping data.

## List of abbreviations

BWA: Burrows-Wheeler Aligner
CDS: coding sequence
FISH: fluorescence *in situ* hybridization
HMM: Hidden Markov Model
Mya: million years ago
MGR: Multiple Genome Rearrangements
SNP: Single Nucleotide Polymorphism

## Acknowledgements

We thank Gayane H. Karagyan and Marine S. Arakelyan for their help with mosquito collection in Armenia and Janet Webster for editing the text.

## Authors’ contributions

Conceptualization: MVS, IVS; experimental design: AAY, MVS, IVS; sampling: GNA, AIV, SMB, MRA, MK, MIG, AVM, BC, SAA, MVS, IVS; molecular-genetic experiments: ANN, DAK, JMH, AIV, AAK, RMB; phylogenetic analyses: AAY; cytogenetic analyses: GNA, VNS, IVS; data analysis: AAY, DAK, IVS. All authors read and approved the final manuscript.

## Availability of data and materials

The data and materials are available from the GenBank repository under the BioProject accession number PRJNA861430.

## Declarations

### Ethics approval and consent to participate

Not applicable.

### Consent for publication

Not applicable.

## References

1. Yurchenko AA, Masri RA, Khrabrova NV, Sibataev AK, Fritz ML, Sharakhova MV: Genomic differentiation and intercontinental population structure of mosquito vectors Culex pipiens pipiens and Culex pipiens molestus. Sci Rep 2020, 10(1):7504.

2. Naumenko AN, Karagodin DA, Yurchenko AA, Moskaev AV, Martin OI, Baricheva EM, Sharakhov IV, Gordeev MI, Sharakhova MV: Chromosome and Genome Divergence between the Cryptic Eurasian Malaria Vector-Species Anopheles messeae and Anopheles daciae. Genes (Basel) 2020, 11(2).

3. Thawornwattana Y, Dalquen D, Yang Z: Coalescent analysis of phylogenomic data confidently resolves the species relationships in the Anopheles gambiae species complex. Mol Biol Evol 2018, 35(10):2512–2527.

4. Lorenz C, Alves JMP, Foster PG, Suesdek L, Sallum MAM: Phylogeny and temporal diversification of mosquitoes (Diptera: Culicidae) with an emphasis on the Neotropical fauna. Syst Entomol 2021, 46(4):798–811.

5. Kamali M, Xia A, Tu Z, Sharakhov IV: A new chromosomal phylogeny supports the repeated origin of vectorial capacity in malaria mosquitoes of the Anopheles gambiae complex. PLoS Pathog 2012, 8(10):e1002960.

6. Aubry F, Dabo S, Manet C, Filipovic I, Rose NH, Miot EF, Martynow D, Baidaliuk A, Merkling SH, Dickson LB et al: Enhanced Zika virus susceptibility of globally invasive Aedes aegypti populations. Science 2020, 370(6519):991–996.

7. Powell JR: Genetic Variation in Insect Vectors: Death of Typology? Insects 2018, 9(4).

8. Rose NH, Sylla M, Badolo A, Lutomiah J, Ayala D, Aribodor OB, Ibe N, Akorli J, Otoo S, Mutebi JP et al: Climate and Urbanization Drive Mosquito Preference for Humans. Curr Biol 2020, 30(18):3570–3579 e3576.

9. Neafsey DE, Waterhouse RM, Abai MR, Aganezov SS, Alekseyev MA, Allen JE, Amon J, Arca B, Arensburger P, Artemov G et al: Mosquito genomics. Highly evolvable malaria vectors: the genomes of 16 Anopheles mosquitoes. Science 2015, 347(6217):1258522.

10. Harbach RE: The classification of genus Anopheles (Diptera: Culicidae): a working hypothesis of phylogenetic relationships. Bull Entomol Res 2004, 94(6):537–553.

11. White GB: Systematic reappraisal of the Anopheles maculipennis complex. Mosquito Systematics 1978, 10:13–44.

12. Krzywinski J, Besansky NJ: Molecular systematics of Anopheles: from subgenera to subpopulations. Annu Rev Entomol 2003, 48:111–139.

13. Sinka ME, Bangs MJ, Manguin S, Coetzee M, Mbogo CM, Hemingway J, Patil AP, Temperley WH, Gething PW, Kabaria CW et al: The dominant Anopheles vectors of human malaria in Africa, Europe and the Middle East: occurrence data, distribution maps and bionomic precis. Parasit Vectors 2010, 3:117.

14. Sinka ME, Rubio-Palis Y, Manguin S, Patil AP, Temperley WH, Gething PW, Van Boeckel T, Kabaria CW, Harbach RE, Hay SI: The dominant Anopheles vectors of human malaria in the Americas: occurrence data, distribution maps and bionomic precis. Parasit Vectors 2010, 3:72.

15. Hackett LW: Malaria in Europe. An ecological study.: Oxford Unoversity Press, London; 1937.

16. Hackett LW, Missiroli A: The varieties of Anopheles maculipennis and their relation to the distribution of malaria in Europe. Riv Malariol 1935, 14:45–109.

17. Kitzmiller JB, Frizzi G, Baker R: Evolution and Speciation within the Maculipennis Complex of the Genus Anopheles. In: Genetics of insect vectors of disease. Edited by Wright JW. Amsterdam-London-New York: Elsevier Publishing Company; 1967: 151–210.

18. Stegniy VN: Reproductive Interrelations of Malarial Mosquitos of the Complex Anopheles-Maculipennis (Diptera, Culicidae). Zool Zh 1980, 59(10):1469–1475.

19. Stegny VN, Sichinava SG, Sipovich NG: A Hybridological Analysis and Biology of the Malarial Mosquitos of the Complex Anopheles-Maculipennis (Diptera, Culicidae) in Western Georgia. Zool Zh 1984, 63(2):300–303.

20. Gutsevich AV, Monchadsky A.S., A.A. S: Fauna of the U.S.S.R. Diptera Volume III No. 4: Moscow, Nauka; 1970.

21. Stegniy VN: Genetic basis of evolution in malaria mosquitoes Anopheles from Maculipennis complex (Diptera, Culicidae). 1. Chromosome-based phylogenetic relationships. Zoologicheskiy Zhurnal 1981, 60(1):69–77.

22. Stegnii VN, Kabanova VM: Cytoecological study of natural populations of malaria mosquitoes on the USSR territory. 1. Isolation of a new species of Anopheles in Maculipennis complex by the cytodiagnostic method. Med Parazitol (Mosk) 1976, 45(2):192–198.

23. Kitzmiller JB: Chromosomal Differences Among Species of Anopheles Mosquitoes. Mosquito Systematics 1977, 9(2):112–122.

24. Stegnii VN, Sharakhova MV: Systemic reorganization of the architechtonics of polytene chromosomes in onto- and phylogenesis of malaria mosquitoes. Structural features regional of chromosomal adhesion to the nuclear membrane. Genetika 1991, 27(5):828–835.

25. Mezzanotte R, Ferrucci L: Recognition of the Sibling Species Anopheles-Atroparvus (Vanthiel) and Anopheles-Labranchiae (Falleroni) (Diptera Culicidae) on the Basis of Q-Banding and C-Banding. Monit Zool Ital 1978, 12(4):211–218.

26. Stegniy VN: Population genetics and evolution of malaria mosquitoes: Tomsk State University Publisher; 1991.

27. Hennig W: Phylogenetic systematics: University of Illinois Press, Urbana, IL; 1966.

28. Stegniy VN: Genetic adaptation and speciation in sibling species of the Eurasian mmaculipennis complex. In: Recent developments in the genetics of insect vectors. Edited by Steiner WM, Tabachnick, W.J., Rai, K.S., Narang, S. Champaign, Illinois; 1982: 454–465.

29. Kitzmiller JB, Baker RH: The salivary chromosomes of Anopheles earlei. Can J Genet Cytol 1965, 7:275–283.

30. Torsvik TH, Cocks RM: Earth history and palaeogeogrphy. Cambridge, UK: Cambridge University Press; 2017.

31. Novikov YM: On the ecology and range of Anopheles beklemishevi (Diptera: Culicidae) with reference to the taxonomy of An. lewisi. J Vector Ecol 2016, 41(2):204–214.

32. Collins FH, Paskewitz SM: A review of the use of ribosomal DNA (rDNA) to differentiate among cryptic Anopheles species. Insect Mol Biol 1996, 5(1):1–9.

33. Porter CH, Collins FH: Phylogeny of nearctic members of the Anopheles maculipennis species group derived from the D2 variable region of 28S ribosomal RNA. Mol Phylogenet Evol 1996, 6(2):178–188.

34. Marinucci M, Romi R, Mancini P, Di Luca M, Severini C: Phylogenetic relationships of seven palearctic members of the maculipennis complex inferred from ITS2 sequence analysis. Insect Mol Biol 1999, 8(4):469–480.

35. Kampen H: The ITS2 ribosomal DNA of Anopheles beklemishevi and further remarks on the phylogenetic relationships within the Anopheles maculipennis group of species (Diptera: Culicidae). Parasitol Res 2005, 97(2):118–128.

36. Djadid ND, Gholizadeh S, Tafsiri E, Romi R, Gordeev M, Zakeri S: Molecular identification of Palearctic members of Anopheles maculipennis in northern Iran. Malar J 2007, 6:6.

37. Gordeev MI, Zvantsov AB, Goriacheva, II, Shaikevich EV, Ezhov MN: [Description of the new species Anopheles artemievi sp.n. (Diptera, *Culicidae*)]. Med Parazitol (Mosk) 2005(2):4–5.

38. Nicolescu G, Linton YM, Vladimirescu A, Howard TM, Harbach RE: Mosquitoes of the Anopheles maculipennis group (Diptera: Culicidae) in Romania, with the discovery and formal recognition of a new species based on molecular and morphological evidence. Bull Entomol Res 2004, 94(6):525–535.

39. Sedaghat MM, Linton YM, Oshaghi MA, Vatandoost H, Harbach RE: The Anopheles maculipennis complex (Diptera: Culicidae) in Iran: molecular characterization and recognition of a new species. Bull Entomol Res 2003, 93(6):527–535.

40. Artemov GN, Bondarenko SM, Naumenko AN, Stegniy VN, Sharakhova MV, Sharakhov IV: Partial-arm translocations in evolution of malaria mosquitoes revealed by high-coverage physical mapping of the Anopheles atroparvus genome. BMC Genomics 2018, 19(1):278.

41. Cahill JA, Soares AER, Green RE, Shapiro B: Inferring species divergence times using pairwise sequential Markovian coalescent modelling and low-coverage genomic data. Philos T R Soc B 2016, 371(1699).

42. Artemov GN, Sharakhova MV, Naumenko AN, Karagodin DA, Baricheva EM, Stegniy VN, Sharakhov IV: A standard photomap of ovarian nurse cell chromosomes in the European malaria vector Anopheles atroparvus. Med Vet Entomol 2015, 29(3):230–237.

43. Bourque G, Pevzner PA: Genome-scale evolution: reconstructing gene orders in the ancestral species. Genome Res 2002, 12(1):26–36.

44. Fontaine MC, Pease JB, Steele A, Waterhouse RM, Neafsey DE, Sharakhov IV, Jiang XF, Hall AB, Catteruccia F, Kakani E et al: Extensive introgression in a malaria vector species complex revealed by phylogenomics. Science 2015, 347(6217):42-+.

45. Small ST, Labbe F, Lobo NF, Koekemoer LL, Sikaala CH, Neafsey DE, Hahn MW, Fontaine MC, Besansky NJ: Radiation with reticulation marks the origin of a major malaria vector. Proc Natl Acad Sci U S A 2020, 117(50):31583–31590.

46. Pringle H: Welcome to Beringia. Science 2014, 343(6174):961–963.

47. Vershinina AO, Heintzman PD, Froese DG, Zazula G, Cassatt-Johnstone M, Dalen L, Der Sarkissian C, Dunn SG, Ermini L, Gamba C et al: Ancient horse genomes reveal the timing and extent of dispersals across the Bering Land Bridge. Mol Ecol 2021.

48. Sun J, Ma PC, Cheng HZ, Wang CZ, Li YL, Cui YQ, Yao HB, Wen SQ, Wei LH: Post-last glacial maximum expansion of Y-chromosome haplogroup C2a-L1373 in northern Asia and its implications for the origin of Native Americans. Am J Phys Anthropol 2021, 174(2):363–374.

49. Lesnek AJ, Briner JP, Lindqvist C, Baichtal JF, Heaton TH: Deglaciation of the Pacific coastal corridor directly preceded the human colonization of the Americas. Sci Adv 2018, 4(5):eaar5040.

50. Ray N, Adams JM: A GIS-based vegetation map of the world at the last glacial maximum (25,000-15,000 BP). Internet Archaelology 2001, 11:1–44.

51. Zvantsov AB, Ejov MN, Artemiev MM: Malaria vectors (Diptera, Culicidae, Anopheles) in CIS countries. In: World Health Organization, Regional Office for Europe, Copenhagen. 2003: 312.

52. Novikov YM, Vaulin OV: Expansion of Anopheles maculipennis s.s. (Diptera: Culicidae) to northeastern Europe and northwestern Asia: causes and consequences. Parasit Vectors 2014, 7:389.

53. Tang C, Davis KE, Delmer C, Yang D, Wills MA: Elevated atmospheric CO2 promoted speciation in mosquitoes (Diptera, Culicidae). Commun Biol 2018, 1:182.

54. Stromberg CA: Evolution of grasses and grassland ecosystems. Annual review of Earth and planetary sciences 2011, 39:517–544.

55. Cantalapiedra JL, Fitzjohn RG, Kuhn TS, Fernandez MH, DeMiguel D, Azanza B, Morales J, Mooers AO: Dietary innovations spurred the diversification of ruminants during the Caenozoic. Proc Biol Sci 2014, 281(1776):20132746.

56. Fabre PH, Hautier L, Dimitrov D, Douzery EJ: A glimpse on the pattern of rodent diversification: a phylogenetic approach. BMC Evol Biol 2012, 12:88.

57. Hanafi-Bojd AA, Vatandoost H, Yaghoobi-Ershadi MR: Climate change and the risk of malaria transmission in Iran. J Med Entomol 2020, 57(1):50–64.

58. Giraldo-Calderon GI, Emrich SJ, MacCallum RM, Maslen G, Dialynas E, Topalis P, Ho N, Gesing S, VectorBase C, Madey G et al: VectorBase: an updated bioinformatics resource for invertebrate vectors and other organisms related with human diseases. Nucleic Acids Res 2015, 43(Database issue):D707–713.

59. Proft J, Maier WA, Kampen H: Identification of six sibling species of the Anopheles maculipennis complex (Diptera: Culicidae) by a polymerase chain reaction assay. Parasitol Res 1999, 85(10):837–843.

60. Bolger AM, Lohse M, Usadel B: Trimmomatic: a flexible trimmer for Illumina sequence data. Bioinformatics 2014, 30(15):2114–2120.

61. Haas BJ, Papanicolaou A, Yassour M, Grabherr M, Blood PD, Bowden J, Couger MB, Eccles D, Li B, Lieber M et al: De novo transcript sequence reconstruction from RNA-seq using the Trinity platform for reference generation and analysis. Nat Protoc 2013, 8(8):1494–1512.

62. TransDecoder (finding coding regions within transcripts)

63. Altschul SF, Madden TL, Schaffer AA, Zhang J, Zhang Z, Miller W, Lipman DJ: Gapped BLAST and PSI-BLAST: a new generation of protein database search programs. Nucleic Acids Res 1997, 25(17):3389–3402.

64. Fu L, Niu B, Zhu Z, Wu S, Li W: CD-HIT: accelerated for clustering the next-generation sequencing data. Bioinformatics 2012, 28(23):3150–3152.

65. Simao FA, Waterhouse RM, Ioannidis P, Kriventseva EV, Zdobnov EM: BUSCO: assessing genome assembly and annotation completeness with single-copy orthologs. Bioinformatics 2015, 31(19):3210–3212.

66. Li H, Durbin R: Fast and accurate short read alignment with Burrows-Wheeler transform. Bioinformatics 2009, 25(14):1754–1760.

67. Li H, Handsaker B, Wysoker A, Fennell T, Ruan J, Homer N, Marth G, Abecasis G, Durbin R, Genome Project Data Processing S: The sequence alignment/map format and SAMtools. Bioinformatics 2009, 25(16):2078–2079.

68. Emms DM, Kelly S: OrthoFinder: solving fundamental biases in whole genome comparisons dramatically improves orthogroup inference accuracy. Genome Biol 2015, 16:157.

69. Loytynoja A: Phylogeny-aware alignment with PRANK. Methods Mol Biol 2014, 1079:155–170.

70. Capella-Gutierrez S, Silla-Martinez JM, Gabaldon T: trimAl: a tool for automated alignment trimming in large-scale phylogenetic analyses. Bioinformatics 2009, 25(15):1972–1973.

71. Darriba D, Taboada GL, Doallo R, Posada D: jModelTest 2: more models, new heuristics and parallel computing. Nat Methods 2012, 9(8):772.

72. Stamatakis A: RAxML version 8: a tool for phylogenetic analysis and post-analysis of large phylogenies. Bioinformatics 2014, 30(9):1312–1313.

73. Kumar S, Stecher G, Tamura K: MEGA7: Molecular Evolutionary Genetics Analysis Version 7.0 for Bigger Datasets. Mol Biol Evol 2016, 33(7):1870–1874.

74. Yang Z: PAML 4: phylogenetic analysis by maximum likelihood. Mol Biol Evol 2007, 24(8):1586–1591.

75. Ward BJ, van Oosterhout C: HYBRIDCHECK: software for the rapid detection, visualization and dating of recombinant regions in genome sequence data. Mol Ecol Resour 2016, 16(2):534–539.

76. Green RE, Krause J, Briggs AW, Maricic T, Stenzel U, Kircher M, Patterson N, Li H, Zhai W, Fritz MH et al: A draft sequence of the Neandertal genome. Science 2010, 328(5979):710–722.

77. Martin SH, Davey JW, Jiggins CD: Evaluating the use of ABBA-BABA statistics to locate introgressed loci. Mol Biol Evol 2015, 32(1):244–257.

78. Ye J, Coulouris G, Zaretskaya I, Cutcutache I, Rozen S, Madden TL: Primer-BLAST: a tool to design target-specific primers for polymerase chain reaction. BMC Bioinformatics 2012, 13:134.

79. Artemov GN, Velichevskaya AI, Bondarenko SM, Karagyan GH, Aghayan SA, Arakelyan MS, Stegniy VN, Sharakhov IV, Sharakhova MV: A standard photomap of the ovarian nurse cell chromosomes for the dominant malaria vector in Europe and Middle East Anopheles sacharovi. Malar J 2018, 17(1):276.

80. Sharakhova MV, Artemov GN, Timoshevskiy VA, Sharakhov IV: Physical Genome Mapping Using Fluorescence In Situ Hybridization with Mosquito Chromosomes. Methods Mol Biol 2019, 1858:177–194.

81. Sharakhov IV: Protocols for cytogenetic mapping of arthropod genomes. Boca Raton, FL.: Taylor & Francis Group, LLC; 2015.

