## Additional file 1 for "Phylogenomics revealed migration routes and adaptive radiation timing of Holarctic malaria vectors of the Maculipennis group"

**Additional file 1. Supplementary online material.**

**Table S1. Statistics of the transcriptome assemblies for species of the Maculipennis group.**

| **Species** | **Total proteins** | **CDS N50, bp** | **Average CDS, bp** | **Total assembled bases** | **BUSCO** |
| --- | --- | --- | --- | --- | --- |
| *An. beklemishevi* | 15135 | 681 | 620.04 | 9384156 | C:28.2%[S:28.0%,D:0.2%],F:21.8%,M:50.0% |
| *An. daciae* Moscow | 21658 | 1014 | 803.73 | 17407329 | C:56.2%[S:54.3%,D:1.9%],F:18.6%,M:25.2% |
| *An. daciae* Tomsk | 21544 | 1212 | 907.56 | 19552251 | C:68.2%[S:66.1%,D:2.1%],F:14.7%,M:17.1% |
| *An. freeborni* | 19832 | 1299 | 986.58 | 19565613 | C:69.4%[S:67.4%,D:2.0%],F:17.4%,M:13.2% |
| *An. labranchiae* | 14404 | 813 | 691.62 | 9962130 | C:36.7%[S:36.5%,D:0.2%],F:21.0%,M:42.3% |
| *An. maculipennis* | 17003 | 930 | 757.8 | 12884655 | C:47.4%[S:47.1%,D:0.3%],F:21.4%,M:31.2% |
| *An. messeae* | 22236 | 1086 | 835.62 | 18580944 | C:62.1%[S:60.0%,D:2.1%],F:17.1%,M:20.8% |
| *An. quadrimaculatus* | 22052 | 1329 | 974.73 | 21495072 | C:73.3%[S:69.7%,D:3.6%],F:12.8%,M:13.9% |
| *An. sacharovi* | 16817 | 1290 | 957.81 | 16107663 | C:64.1%[S:63.8%,D:0.3%],F:17.2%,M:18.7% |

**Table 2. Genes of *An. atroparvus* used as markers for detection of interspecies chromosome rearrangements on the X chromosome in five species of the Maculipennis group.** The distance between genes was calculated as the number of nucleotides from the start to the end of a gene on the *An. atroparvus* genome map.

|  | Gene ID | Gene start (bp) | Gene end (bp) | Distance from the previous gene (bp) | Supercontig | Chromosome band |
| --- | --- | --- | --- | --- | --- | --- |
| 1 | AATE010696 | 18023 | 22191 |  | KI421896 | 1A |
| 2 | AATE001169 | 1162806 | 1170169 | 1140615 | KI421896 | 1A |
| 3 | AATE017741 | 1960425 | 1963110 | 790256 | KI421896 | 1B |
| 4 | AATE010870 | 2927563 | 2928974 | 964453 | KI421896 | 1C |
| 5 | AATE017428 | 3964271 | 3982124 | 1035297 | KI421896 | 2A |
| 6 | AATE012020 | 4602182 | 4608814 | 620058 | KI421895 | 2A |
| 7 | AATE015765 | 5527302 | 5531343 | 918488 | KI421895 | 2B |
| 8 | AATE000795 | 6504256 | 6513151 | 972913 | KI421895 | 2B |
| 9 | AATE002125 | 7208125 | 7217523 | 694974 | KI421895 | 2C |
| 10 | AATE004111 | 8538061 | 8542551 | 1320538 | KI421895 | 2C |
| 11 | AATE001403 | 8751285 | 8754177 | 208734 | KI421895 | 2C |
| 12 | AATE018270 | 9855878 | 9857040 | 1101701 | KI421907 | 3B |
| 13 | AATE017493 | 10389518 | 10398371 | 532478 | KI421907 | 3B |
| 14 | AATE021170 | 11446721 | 11457749 | 1048350 | KI421907 | 3C |
| 15 | AATE018218 | 11958281 | 11959687 | 500532 | KI421898 | 3C |
| 16 | AATE005236 | 12437092 | 12444184 | 477405 | KI421898 | 3C |
| 17 | AATE010434 | 13484949 | 13487325 | 1040765 | KI421898 | 4A |
| 18 | AATE009010 | 14525064 | 14530105 | 1037739 | KI421898 | 4A |
| 19 | AATE018661 | 15491805 | 15495758 | 961700 | KI421898 | 4A |
| 20 | AATE010210 | 15918625 | 15921507 | 422867 | KI421919 | 4B |
| 21 | AATE016157 | 16324241 | 16326290 | 402734 | KI421920 | 4B |

**Table S3. Mosquito species and sampling sites.**

| **Species** | **Country** | **Region** | **Location** | **GPS coordinates** | **Date of collection** |
| --- | --- | --- | --- | --- | --- |
| *An. artemievi* | Kyrgyzstan | Osh | Kydyrsha and Kyzyl Shark | 40.726613, 72.957595; 40.725281, 72.932879 | 08/22/2006 |
| *An. beklemishevi* | Russia | Tomsk | Chainsk | 57.931778, 82.596862 | 08/22/2016 |
| *An. daciae* Moscow | Russia | Moscow | Novokosino | 55.734170, 37.843453 | 08/28/2015 |
| *An. daciae* Tomsk | Russia | Tomsk | Kandinka | 56.292605, 84.806183 | 07/08/2015 |
| *An. freeborni* | USA | California | Marysville | Unknown (colony) | Established in 1943 |
| *An. quadrimaculatus* | USA | Florida | Orlando | Unknown (colony) | Established in 1939 |
| *An. labranchiae* | Italy | Tuscany | Princhipina terra | 42.724717, 11.041467 | 08/02/2015 |
| *An. maculipennis* | Italy | Lacio | Rieti | 42.404833, 12.829150 | 08/05/2015 |
| *An. martinius* | Kazakhstan | Kysyl-Orda | Kyzil-Orda | 44.852417, 65.118217 | 08/23/2005 |
| *An. melanoon* | Georgia, Abkhazia | Gudauta | Primorskiy | 43.09351, 40.69351 | 07/04/2017 |
| *An. messeae* | Russia | Moscow | Novokosino | 55.734170, 37.843453 | 08/28/2015 |
| *An. persiensis* | Iran | Mazandaran | Alendan, Andar-Gholi, Chader-Mahaleh, Chalmardi | 36.221983, 53.433833; 36.339467, 52.883750; 36.513650, 52.345100; 36.561389, 53.392500 | 09/07/2017  09/06/2017  05/19/2017 |
| *An. sacharovi* | Armenia | Ararat | Araksavan | 39.784329, 45.229700 | 09/02/2016 |
